# A parametric logistic equation with light flux and medium concentration for cultivation planning of microalgae

**DOI:** 10.1101/2022.02.28.482388

**Authors:** Kazuki Kambe, Yasutaka Hirokawa, Asuka Koshi, Yutaka Hori

## Abstract

Microalgae are considered to be promising producers of bioactive chemicals, feeds, and fuels from carbon dioxide by photosynthesis. Thus, the prediction of microalgal growth profiles is important for the planning of cost-effective and sustainable cultivation-harvest cycles. This paper proposes a mathematical model capable of predicting the effect of light flux into culture and medium concentration on the growth profiles of microalgae by incorporating these growth-limiting factors into the logistic equation. The specific form of the equation is derived based on the experimentally measured growth profiles of *Monoraphidium* sp., a microalgal strain isolated by the authors, under 16 conditions consisting of the combinations of incident light fluxes into culture and initial medium concentrations. Using a cross-validation method, it is shown that the proposed model has the ability to predict necessary incident light flux into culture and initial medium concentration for harvesting target biomass at target time. Finally, the model-guided cultivation planning is performed and is evaluated by comparing the result with the experimental data.

## 1 Introduction

Photosynthesis is a beneficial reaction that reconverts atmospheric carbon dioxide produced by the consumption of fossil resources into organic carbon. It is one of the desirable solutions for a sustainable future to substitute photosynthetic products for fossil resources. Microalgae, which grow faster and show higher carbon fixation rates than higher plants, are excellent candidates for carbon neutral producer. Some species of microalgae have been used as live feed for fish larvae and are expected as alternative feedstocks for livestock [1] and aquaculture [2] because of their high nutritional quality. Other species of microalgae were reported as superior producers of bioactive chemicals [3], biofuels [4,5], and biohydrogen [6], which would potentially revolutionize the production of cosmetics, health foods, and fossil-based energy. Among them, *Monoraphidium* genera classified in Selenastraceae are oleaginous microalgae showing high lipid contents and were reported as promising hosts for lipid biofuel production [5,7–10]. Moreover, recent studies showed that *Monoraphidium* could grow robustly even in wastewater, suggesting that *Monoraphidium* cultivation would become a simultaneous solution for bioremediation and biorefinery [11, 12]. To date, optimization of microalgal cultivation has been studied by computational and experimental approaches [13–15]. In these types of optimization, it is a common procedure to optimize the titer of microalgae in a single culture harvest. However, a major challenge in practical algal cultivation lies in the discovery of cultivation conditions that enable continuously stable and cost-effective production over multiple cultivation-harvest cycles. Thus, a model-guided approach to the design of a cultivation plan for long-term cultivation is desirable. In particular, development of a mathematical model capable of predicting the number of cells and harvest time in response to various cultivation conditions will be a key to finding the conditions for sustainable cultivation.

Many existing mathematical models predict the growth rate in response to various environmental factors during the cultivation [16, 17]. Examples include the Monod’s growth model describing the effect of specific nutrients [18] and the models of temperaturedependent cell growth [14, 15, 19]. The effect of light intensity on the growth profile was also modeled in various ways depending on the state of the cell culture [17]. Although the growth rate is affected by numerous factors including nutrients concentration, light intensity, temperature, carbon dioxide concentration, and toxic byproducts in the medium [20], these models consider the effect of only a few factors based on the assumption that cell growth is ultimately limited by those factors. In contrast to these models, the logistic equation incorporates the bulk effect of multiple growth limiting factors such as medium concentration and toxic byproducts into a single parameter called the carrying capacity of the environment [21]. Thus, the equation was used to fit the growth profiles of a wide range of microorganisms [22–26]. However, since it does not explicitly consider the counteraction of the biomass production to the change of environmental factors, the original logistic equation cannot directly be used for the exploration of the growth conditions in the cultivation planning. To overcome the limitations of these models, a recent study proposed a hybrid Logistic-Monod model [27]. This model explicitly takes the medium concentration into the variable while the other self-limiting factors are considered in the logistic-type equation. In particular, this model explicitly captures the dynamic interplay of the biomass production and the medium consumption using ordinary differential equations (ODEs) to enable the exploration of cultivation conditions. This idea of modeling the complex interplay of biomass production and environmental factors can be used to build a more advanced hybrid model that includes other important factors for algal cultivation such as light intensity.

This paper proposes an ODE model of the growth profile of microalgae to enable cultivation planning by model-based exploration of the cultivation parameter space, as shown in Fig. 1 (A). Specifically, the proposed model extends the logistic equation in a way that explicitly incorporates the dynamic environmental factors such as light flux being available for a single cell and medium concentration, enabling to find a cultivation condition that satisfies various constraints of practical cultivation process such as target biomass and target time. The specific form of the extended logistic equation and its parameters were determined from experimentally measured growth profiles of an originally isolated *Monoraphidium* sp. under different light fluxes into culture and medium concentrations. These experimental data were further used for the cross-validation of the proposed model. The cross-validation result showed that the model could replicate various features of the growth profiles, including the peak biomass and its timing, for different cultivation conditions. Finally, we showcase a procedure of cultivation planning, where the initial medium concentration and the incident light flux into culture are explored to achieve predefined target biomass and target time based on the simulations and some analytic relation of the proposed model.

**Figure 1:**
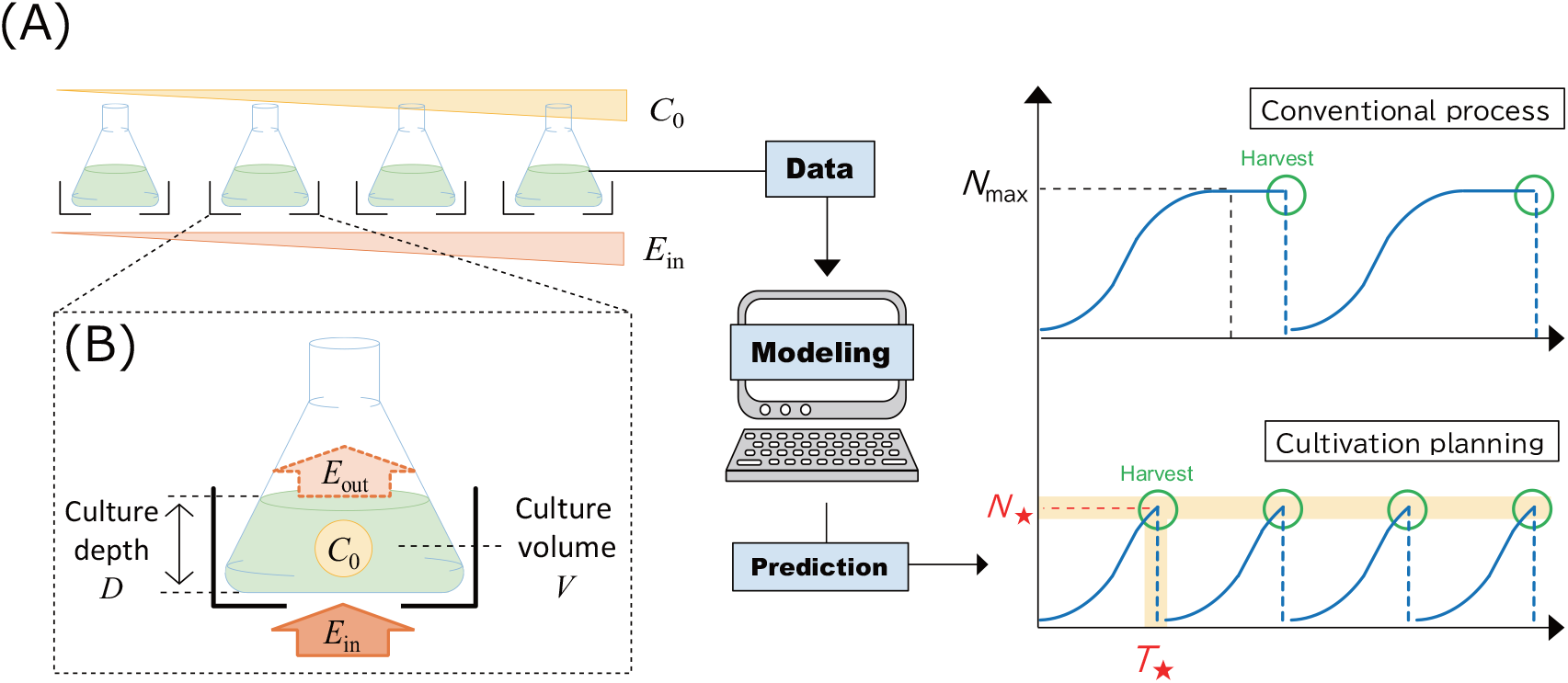
(A) Overview of cultivation planning. The symbols (*C*_0_, *E*_in_, *N*_max_, *N*_⋆_ and *T*_⋆_) are the initial medium concentration, the incident light flux into culture, the maximum biomass, the target biomass, and the target time, respectively. (B) Enlarged view of experimental setup. The symbol *E*_out_ is the transmitted light flux thorough culture.

## 2 Experimental conditions and overview of the proposed model

The goal of cultivation planning is to find parameters of cultivation such as the initial medium concentration *C*_0_ and the incident light flux into culture *E*_in_ that achieve a target biomass *N*_⋆_ at target time *T*_⋆_. To this end, we build a mathematical model that can predict the dynamic biomass *N* in response to various cultivation parameters as shown in Fig. 1 (A). In this section, we first introduce experimental conditions for building the proposed model, and then outline the overview for the proposed model.

### 2.1 Experimental conditions

Microalgae isolated from a freshwater pond in Yoshikawa, Saitama, Japan was named ACCB1808. The 18S ribosomal DNA sequence of ACCB1808 was 99.3 %, 99.1 %, and 99.1 % identical to that of *Monoraphidium* sp. LB59, *M. subclavatum* FBCC-A409, and *Monoraphidium* sp. HDMA-11, respectively (Fig. S.1). This result indicated that ACCB1808 belonged to the genus of *Monoraphidium*.

To measure the growth profile for modeling, ACCB1808 cultures were incubated in 300-mL flasks containing 200 mL of modified BG11 medium. The details of experimental conditions including the medium composition were summarized in Appendix A. Flasks containing culture were directly placed onto white LED light whose intensity measured as photosynthetic photon flux density (PPFD) was 1034 µE · m^−2^ · s^−1^. To strictly regulate the incident light flux into culture, black drawing paper with a 3-cm radius hole (area of 2.826×10^−3^ m^2^) was inserted between the flasks and the LED light. The flasks were placed in an enclosure made with the same drawing paper as shown in Fig. 1 (B). The incident light intensities to flasks measured as PPFD were adjusted to 1034, 386.7, 184.8, and 96.8 µE · m^−2^ · s^−1^ by inserting sheets of papers between the flasks and the LED light. Owing to strict regulation of the illuminated area (2.826 × 10^−3^ m^2^), the incident light fluxes into culture *E*_in_ were calculated as 2.92, 1.09, 0.521, and 0.274 µE · s^−1^. The concentration of modified BG11 medium *C* was defined as 1, and the initial medium concentrations *C*_0_ were adjusted to 0.5, 0.25, and 0.125 by dilution. Although the medium is composed of different nutrients consumed at various rates upon growth, we here assume that there is a rate-limiting nutrient species and use the single variable *C* to capture the growth limiting effect by that nutrient. ACCB1808 cultures were incubated under 16 conditions consisting of the combinations of four patterns of initial medium concentration *C*_0_ and four patterns of incident light fluxes into culture *E*_in_ as shown in Fig. 2 (A). These experimental data were used for building and evaluating the proposed model in the following sections.

**Figure 2:**
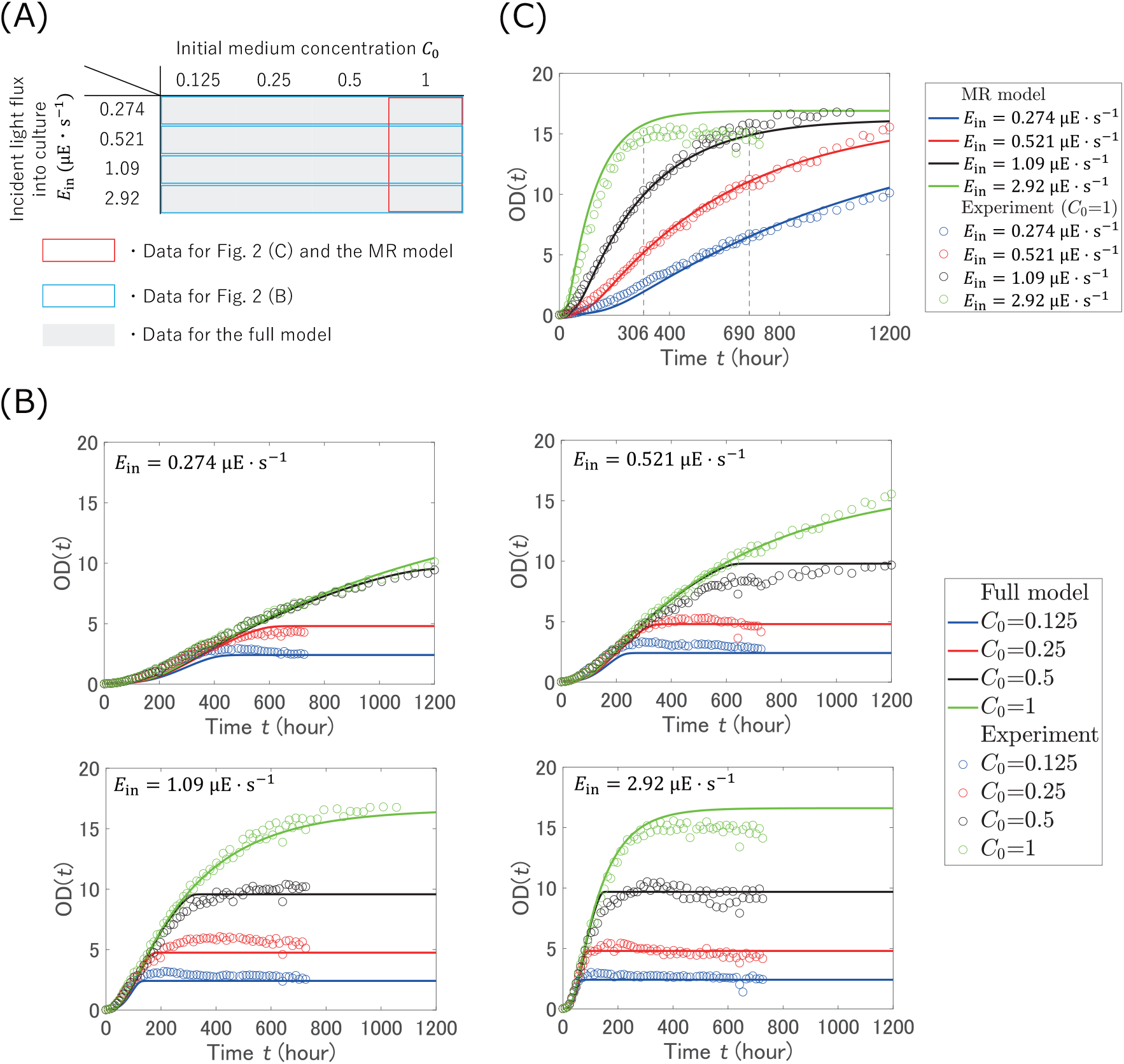
(A) Experimental conditions combining initial medium concentrations *C*_0_ and light fluxes into culture *E*_in_. (B) Time series data for different initial medium concentrations *C*_0_ and incident light flux into culture *E*_in_. Circles and solid lines show experimental data and predicted results of the full model described in Section 4, respectively. (C) Time series data for the four incident light fluxes into culture *E*_in_ with *C*_0_ = 1. Circles and solid lines show experimental data (repeat of the data in panel (B)) and predicted results of the MR model described in Section 3, respectively.

### 2.2 Overview of the proposed model

The growth profiles of experimental data shown in Fig. 2 (B) indicated that the growth rate and the biomass accumulation were dependent on the incident light flux into culture *E*_in_ and the initial medium concentration *C*_0_, respectively. In initial growth phase, the biomass *N* increased exponentially. Next, after following linear increase, the growth rate gradually decreased and leveled off. These phase transitions were possibly caused by photoinhibition, insufficient light absorption at high biomass, depletion of the medium concentration, and increase of inhibitory substances [28, 29]. The state of culture used for subculture often affected the initial growth rate in the next cultivation. Specifically, the culture state before stationary phase was preferable for use in subculture. To harvest the culture in the preferable state, it is important to predict not only a target biomass *N*_⋆_ but also the target time *T*_⋆_ at which the biomass reaches *N*_⋆_. Moreover, for practical operation, the target values (*N*_⋆_ and *T*_⋆_) have to be decided from various viewpoints such as cost effectiveness and the schedule of operators. A mathematical model would be a desirable tool for exploring cultivation conditions satisfying these target values.

In what follows, we build a mathematical model capable of predicting the growth kinetics of ACCB1808 culture for various cultivation conditions. In particular, we are interested in the growth kinetics in response to light flux into culture *E*_in_ and medium concentration *C* since these parameters largely affect the growth profile as shown in Fig. 2(B). Since the increase of biomass decreases available light flux for a single cell even if light flux into culture *E*_in_ is constant during cultivation, available light flux for a single cell is one of the key variables for the prediction of the dynamic biomass change. In this paper, light flux being available for a single cell is called light flux per cell *L*. The effect of light flux per cell on the growth kinetics will be discussed in Section 3.

We develop an ordinary differential equation (ODE) model of the growth kinetics based on the logistic equation [21]:

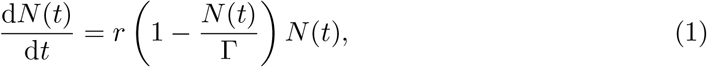

where *N* (*t*) is the biomass at time *t, r* is the maximum specific growth rate, and Γ is the carrying capacity of the environment, or the maximum achievable population size. The first term of the logistic equation represents the growth due to proliferation, and the second term collectively accounts for growth-limiting factors such as toxic byproducts and reactive oxygen species.

Unlike the original logistic equation [21], we assume that the maximum specific growth rate *r* is dependent on the light flux per cell *L*, and the medium concentration *C*, that is, *r* := *r*(*L, C*). As discussed later in detail, *L* and *C* are subject to dynamic change since these two variables are affected by biomass *N* (*t*). The combined effect of these factors is incorporated into the maximum specific growth rate *r*(*L, C*) by

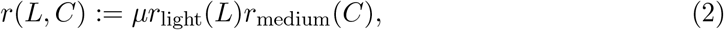

where *µ* is a constant, and *r*_light_(*L*) and *r*_medium_(*C*) are functions of light flux per cell and medium concentration. These functions take values between 0 and 1. The specific forms of these functions are defined in Section 3 and 4. The functions *r*_light_(*L*) and *r*_medium_(*C*) represent the growth-limiting effect due to insufficient absorbance of light flux per cell and insufficient nutrients, respectively. In other words, *r*(*L, C*) ≃ *µ* when the light flux per cell *L* and the medium concentration *C* are sufficiently high such that *r*_light_(*L*) ≃ 1, and *r*_medium_(*C*) ≃ 1, while *r*(*L, C*) ≃ 0 when *L* or *C* is close to 0.

## 3 Logistic equation with light flux per cell

In the previous section, the maximum specific growth rate *r* is defined by the functions of the medium concentration *C* and of the light flux per cell *L*. When the medium concentration *C* is sufficiently high, the maximum specific growth rate *r* is only dependent on the light flux per cell *L*. In this section, we consider this special case and formulate a logistic equation that incorporates the effect of the light flux per cell *L*, which we call the medium-rich model or the MR model for shorthand.

### 3.1 Modeling of the MR model

We incorporate the effect of the light flux per cell *L* on the growth kinetics by defining

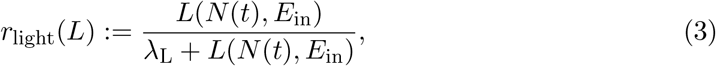

where *λ*_L_ is a half-velocity constant satisfying *r*_light_(*λ*_L_) = 1/2. The light flux per cell *L*(*N* (*t*), *E*_in_) is dependent on the incident light flux into culture *E*_in_ and the biomass *N* (*t*). The light flux absorbed by the culture is expressed by *E*_in_ − *E*_out_ where *E*_out_ is the transmitted light flux through culture as illustrated in Fig. 1 (B) (see details about *E*_out_ in Appendix B). Thus, the absorbed light flux per cell *L*(*N* (*t*), *E*_in_) is written as (*E*_in_ − *E*_out_)*/N* (*t*) by assuming that light flux is uniformly absorbed by cells in culture. Moreover, the attenuation of transmitted light in the culture obeys Lambert-Beer’s law [30], which states the exponential decrease of light flux with the path length of light flux and the concentration of the solution (biomass concentration) (see details in Appendix C). Thus, the absorbed light flux per cell *L*(*N* (*t*), *E*_in_) is

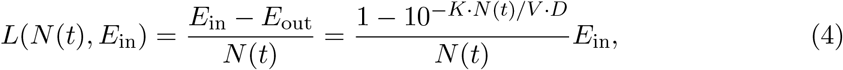

where *K* is the cell-specific extinction coefficient, *D* is the culture depth, and *V* is the culture volume as illustrated in Fig. 1 (B). It should be noted that, in general, the profile of the growth rate functions of light flux per cell could be sigmoidal at low light flux per cell *L* due to the minimum light flux required for cell growth, and be decreasing at high light flux due to photoinhibition [28]. Thus, Eq. (3) is an approximation model that only captures the increasing and the saturation phase of the growth rate in the mild light flux condition, where the cultivation is mainly performed.

When the nutrients in the medium are sufficient and are not rate-limiting factors, *r*_medium_(*C*) = 1 holds. Thus, it follows that

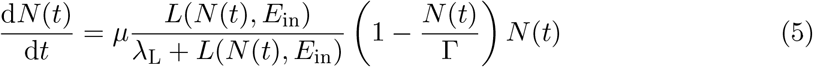

by substituting Eq. (3) into Eq. (1). In what follows, Eq. (5) is called the MR model. The MR model allows us to assess some of the parameters of Eqs. (1) and (2) by using experimental data taken under the conditions with sufficient medium concentration as shown in the next subsection. The other parameters that appear in *r*_medium_(*C*) can then be assessed by subsequent experiments, which will be discussed in Section 4. The two-step parameter evaluation helps avoid overfitting to a single experimental data.

### 3.2 Parameter evaluation of the MR model

Experiments were conducted to assess the parameters *K, µ, λ*_L_, and Γ of the MR model (Eq. (5)). Firstly, the extinction coefficient *K* being specific to ACCB1808 was evaluated by the method of Masuda *et al*. [30]. PPFD was measured in ACCB1808 culture with various biomass concentrations and culture depths (see details in Appendix C). The relative logarithmic PPFD was negatively correlated with biomass concentrations and culture depths, indicating that Lambert-Beer’s law was obeyed in ACCB1808 culture. When the units of culture depth and biomass concentration were defined as “cm” and “cell · mL^−1^”, the extinction coefficient *K* was evaluated as *K* = 5.1 × 10^−9^ mL · cm^−1^ · cell^−1^.

Using this extinction coefficient, we further performed evaluation of *µ, λ*_L_ and Γ in the MR model (Eq. (5)) based on the growth profile of ACCB1808 culture (see details about cultivation conditions in Appendix A). Experiments were conducted under a total of 16 different conditions as shown in Fig. 2 (A), where the growth curves were obtained for the combinations of four different light fluxes into culture, 0.274, 0.521, 1.09 and 2.92 µE · s^−1^, and four initial medium concentrations, 0.125, 0.25, 0.5 and 1. Optical density at 730 nm (OD) was measured as a proxy of biomass *N* (*t*) (see details in Appendix C). The parameters were then fitted to the four growth kinetics with the sufficient medium concentration, *i*.*e*., *C*_0_ = 1 in Fig. 2 (C). Specifically, the culture volume *V*, the culture depth *D*, the initial optical density OD_0_, and the evaluated extinction coefficient *K* were set in Eq. (5) as shown in Table 1. When 200 mL culture was put into 300-mL flask, culture depth corresponded 3.7 cm. The culture depth *D* and the culture volume *V* were assumed to remain constant during cultivation. For *E*_in_ = 1.09 and 2.92 µE · s^−1^, time series data were fitted only for the first 690 and 306 hour, respectively, since the medium concentration could become a rate-limiting factor, violating the assumption of the MR model, when the culture reached stationary phase.

**Table 1:** Initial values and assessed extinction coefficient *K*.

| Parameter | Unit | Value |
| --- | --- | --- |
| $V$ | mL | 200 |
| $D$ | cm | 3.7 |
| $OD_0$ | - | 0.025 |
| $K$ | $\text{mL} \cdot \text{cm}^{-1} \cdot \text{cell}^{-1}$ | $5.1 \times 10^{-9}$ |

The assessed parameters are shown in Table 2. This result implied that the maximum specific growth rate *r*(*L, C*) ≃ *µr*_light_(*L*) was between 0 and 0.194 hour^−1^ depending on the light flux per cell *L* when the medium concentration was high, *i*.*e*., *r*_medium_(*C*) ≃ 1. The carrying capacity Γ was evaluated as 99.9 × 10^9^ cell. The biomass concentration 99.9 × 10^9^/200 cell · mL^−1^ can be converted into OD of 16.6. This indicated that the value of OD would never exceed 16.6 regardless of the medium concentration.

**Table 2:** List of evaluated parameters of the proposed model.

| Parameter | Unit | Assessed value |
| --- | --- | --- |
| $\mu$ | $\text{hour}^{-1}$ | 0.194 |
| $\lambda_L$ | $\mu\text{E} \cdot \text{s}^{-1} \cdot \text{cell}^{-1}$ | $1.90 \times 10^{-6}$ |
| $\Gamma$ | cell | $99.9 \times 10^9$ |
| $\xi_C$ | - | 0.012 |
| $\alpha$ | $\text{cell}^{-1}$ | $8.7 \times 10^{-12}$ |

To evaluate the predictive ability of the MR model, the predicted results of the MR model were further compared with the experimental data using leave-one-out cross-validation (LOOCV) [31], where three of the four experimental conditions were grouped together for parameter evaluation and the other was used for prediction. Specifically, the parameters (*µ, λ*_L_ and Γ) were fitted to the three growth curves with *C*_0_ = 1. Then, the model was used to predict the growth kinetics as shown in Fig. 2 (C). Figure 2 (C) shows that the MR model was capable of predicting the growth kinetics before reaching stationary phase, where the decrease in medium concentration had little effect on cell growth.

In the next section, the MR model is used to build a full model that captures the effect of both light flux per cell *L* and medium concentration *C* on cell growth.

## 4 Logistic equation with light flux per cell and medium concentration

A standing assumption of the MR model is that the medium concentration *C*(*t*) is sufficiently high so that the maximum specific growth rate *r* is independent of *C*(*t*). In this section, we will extend the MR model (Eq. (5)) to remove this assumption and explicitly incorporate the effect of the medium concentration on the maximum specific growth rate.

### 4.1 Modeling of logistic equation with light flux and medium concentration

The experimental data in Fig. 2 (B) suggest that the medium concentration *C*(*t*) does not affect the growth profile just before the stationary phase is reached. Based on this observation, we incorporate the rate-limiting effect of the medium concentration using the Monod-type model [18]:

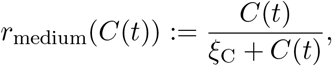

where *ξ*_C_ is a half-velocity constant. We assume that nutrients in the medium are mainly used for cell growth, and the consumption rate is proportional to the growth rate d*N* (*t*)/d*t*. Consequently, the mathematical model that incorporates the effect of both the light flux per cell *L*(*N* (*t*), *E*_in_) and the medium concentration *C*(*t*) is obtained as

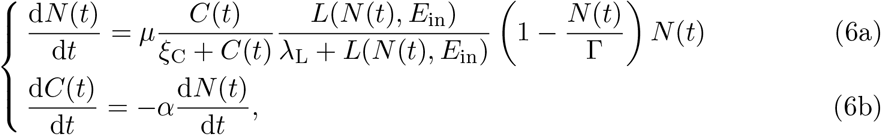

where *α* is a parameter representing consumption of the medium concentration per a unit increase of the biomass, and *L*(*N* (*t*), *E*_in_) is defined by Eq. (4). Eq. (6) is called the full model in the following sections. Defining the initial biomass *N* (0) by *N*_0_, Eq. (6) can equivalently be expressed as

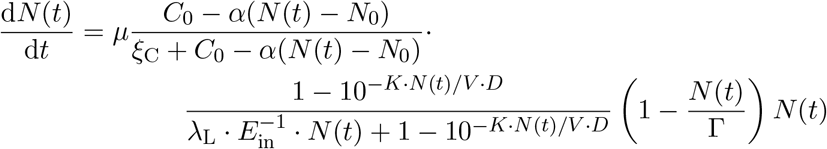

since Eq. (6b) implies

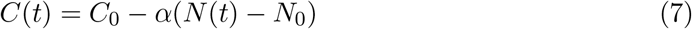

and *E*_in_ *>* 0 after starting cultivation.

It should be noted that the parameters *K, µ, λ*_L_, and Γ are assessed in the MR model as described in Section 3.2. The other parameters *ξ*_C_ and *α* can be assessed using the 16 experimental data for the different incident light fluxes into culture *E*_in_ and the initial medium concentrations *C*_0_ in Fig. 2 (B).

### 4.2 Parameter assessment in the full model

The parameters *ξ*_C_ and *α* in the full model (Eq. (6)) were assessed using the 16 experimental data in Fig. 2 (B). Specifically, the initial parameters, *V, D*, and OD_0_, and the assessed extinction coefficient *K* in Table 1 were used. The parameters *µ, λ*_L_, and Γ obtained using the MR model in Section 3.2 were set (see Table 2). Then, the least square solution was searched for *ξ*_C_ and *α*. The evaluated parameters are shown in Table 2.

The generalizability of the full model was also evaluated by LOOCV [31] using the experimental conditions matrix in Fig. 2 (A). Specifically, for each combination of the light flux into culture *E*_in_ and the initial medium concentration *C*_0_, the parameters *ξ*_C_ and *α* were assessed with the other 15 experimental data. Then, the model was used to predict the growth kinetics as shown in Fig. 2 (B). The simulated growth kinetics showed agreement with the experimental data in that the dynamic of the growth rate was dependent on the light flux into culture *E*_in_ before reaching stationary phase, while the maximum biomass was dependent on the initial medium concentration *C*_0_. The result of LOOCV in Fig. 2 (B) also suggests that the model can predict the maximum biomass *N*_max_ or its corresponding maximum OD. These results will be more quantitatively evaluated in the next section along with the demonstration of cultivation planning.

## 5 Demonstration of cultivation planning

The goal of cultivation planning is to find the initial medium concentration 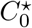 and the incident light flux into culture 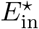 to achieve a predefined target biomass *N*⋆ at target time 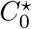 In a typical cultivation process, cells are harvested before reaching the stationary phase to avoid the carry-over of potentially toxic byproducts in subculture. Thus, 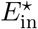 and 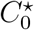 should be planned so that biomass reaches *N*_⋆_ at *T*_⋆_.

### 5.1 Prediction of initial medium concentration *C*^⋆^ for target biomass *N*_⋆_

In a typical cultivation cycle, culture is harvested before reaching stationary phase to maintain a preferable culture state in subculture. In other words, target biomass *N*_⋆_ should be set less than the maximum biomass at stationary phase *N*_max_, *e*.*g*., *N*_⋆_ = 0.8*N*_max_. Thus, prediction of the maximum biomass *N*_max_ or its corresponding maximum OD in response to cultivation conditions such as incident light flux into culture *E*_in_ and initial medium concentration *C*_0_ is important in cultivation planning. In theory, *N*_max_ is the biomass at steady state, at which d*N* (*t*)/d*t* = 0. The steady state is achieved either when the biomass *N* (*t*) reaches the carrying capacity Γ, *i*.*e*., *N* (*t*) = Γ, or the medium concentration is depleted, *i*.*e*., *C*(*t*) = 0. Thus, *N*_max_ can be analytically calculated from Eqs. (6) and (7) as

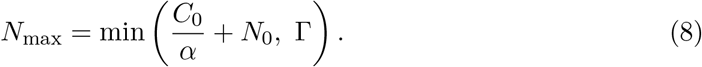

It should be noted that the biomass *N* (*t*) can be converted to OD by

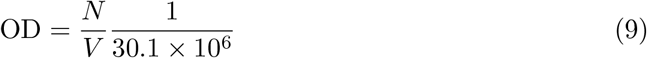

and dry weight per OD is 0.213 mg · mL−1. Eq. (8) implies that the maximum biomass *N*_max_, or its corresponding maximum OD, is obtained from an initial medium concentration *C*_0_ when *C*_0_ *< α*(Γ − *N*_0_). This analytic solution provides a crude estimation of biomass at stationary phase for a given initial medium concentration.

Figure 3 shows the maximum OD predicted from Eqs. (8) and (9), and measured by the experiments in Fig. 2 (B), where the parameters *α* and Γ in Table 2 and the initial OD_0_ in Table 1 corresponding the initial biomass *N*_0_ were used for calculation. The results show that the maximum OD is determined from the initial medium concentrations *C*_0_ and is independent of the incident light fluxes into culture *E*_in_. Thus, a predefined target biomass *N*_⋆_ or its corresponding target OD for cultivation planning is obtained by simply selecting the initial medium concentration 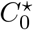, which can be calculated from Eq. (8). Once the initial medium concentration 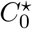 is fixed, the time when biomass reaches *N*_⋆_ can be adjusted by incident light flux into culture *E*_in_. In the next subsection, we will give a demonstration to select the incident light flux into culture 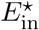 for target time *T*_⋆_.

**Figure 3:**
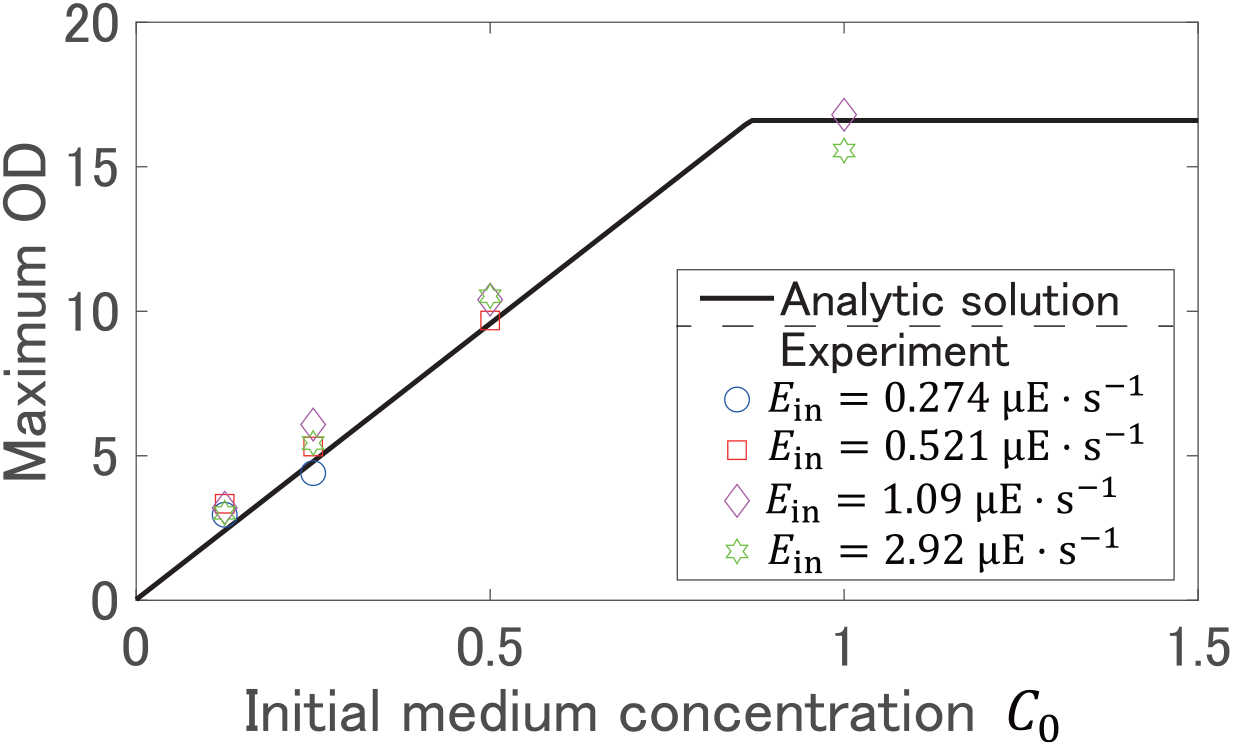
Maximum OD calculated from the analytic solution (Eqs. (8) and (9)) and measured by experiments in Fig. 2 (B).

### 5.2 Prediction of incident light flux into culture 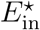 for target time *T*_⋆_

Once the initial medium concentration 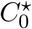 is selected based on Eq. (8), the next goal is to seek the incident light flux into culture 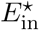 that achieves target biomass *N*_⋆_ at target time *T*_⋆_. The incident light flux into culture 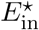 is explored by running simulations of the full model (Eq. (6)). Since the target biomass *N*_⋆_ is often set less than the maximum biomass at stationary phase in practical cultivation, let us suppose, for example, that the target biomass *N*_⋆_ is around 70–90% of the maximum biomass. Then, the expected harvest time, at which the biomass reaches the target biomass can be computed from the growth kinetics simulated by Eq. (6) for each *E*_in_.

Figure 4 shows the time spans in which the biomass reaches 70–90% of its maximum value for different incident light fluxes into culture *E*_in_. In Fig. 4, *T*_1_ and *T*_2_ represent the simulated lower and upper bound of harvest time, at which biomass reaches 0.7*N*_max_ and 0.9*N*_max_, respectively. The parameters in Tables 1 and 2 were used for these simulations of the full model (Eq. (6)). The experimental data in Fig. 4 is obtained from the time-series data in Fig. 2 (B).

**Figure 4:**
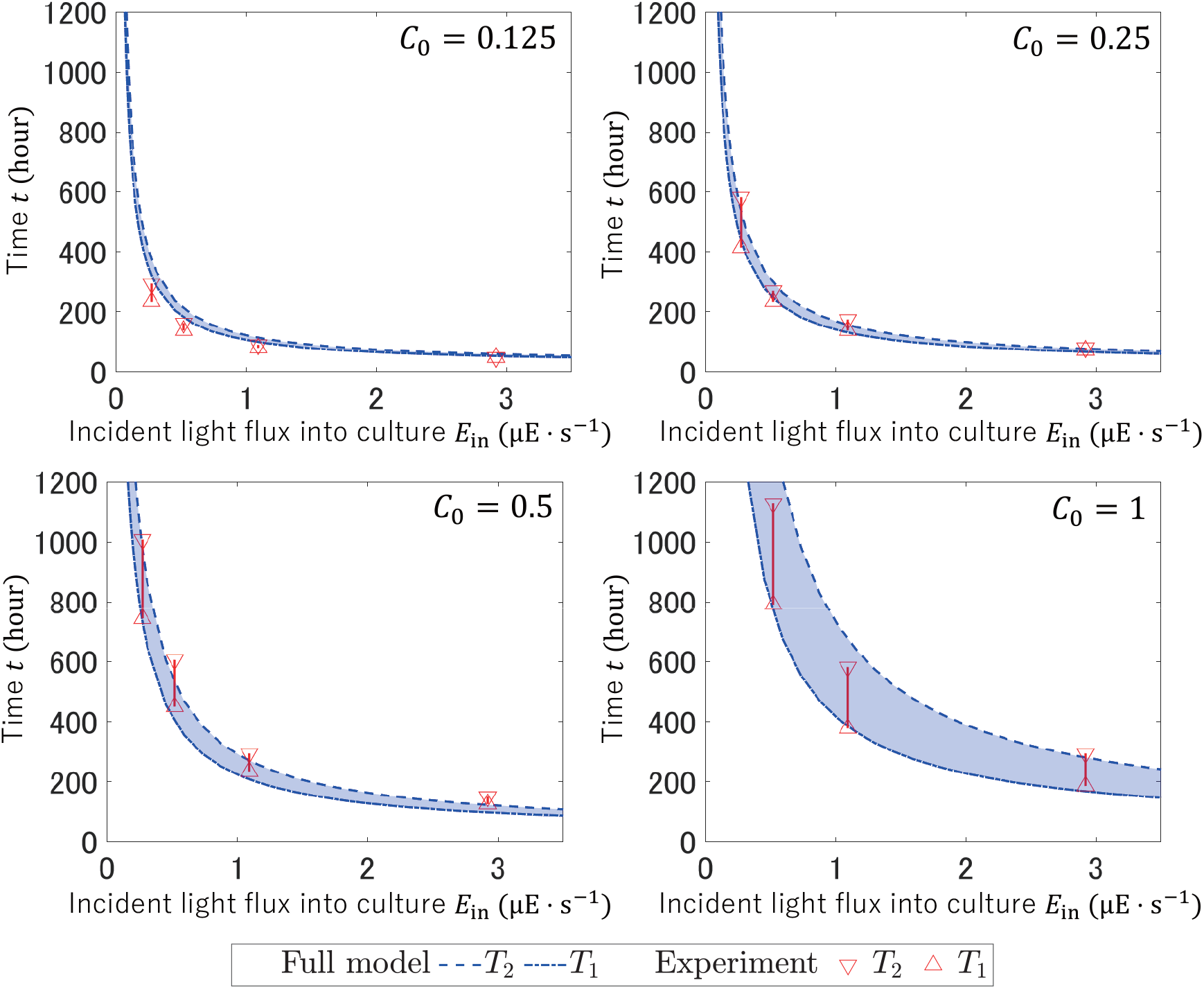
Span of target time obtained by simulations of the full model (Eq. (6)) and measured by experiments in Fig. 2 (B).

Figure 4 shows agreement of the computationally predicted time spans with the experimental data. Thus, the incident light flux into culture 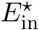 that enables to harvest target biomass at target time can be computationally determined based on the full model (Eq. (6)). Thus, combining with the prediction of the initial medium concentration 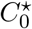 in Section 5.1, one can find cultivation conditions (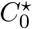 and 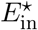) satisfying the predefined constraints (*N*⋆ and *T*⋆), fulfilling the goal of cultivation planning.

## 6 Conclusion

Microalgae are photosynthetic organisms that have high potential as carbon neutral producer and alternative feedstocks for livestock [1] and aquaculture [2]. Prediction of biomass in cultivation of microalgae is a difficult task due to the complex interplay of growth conditions such as light flux, medium concentration, and temperature [20]. As a result, experimental conditions for harvesting target biomass at target time, *i*.*e*., cultivation plans, are often sought empirically by operators.

This paper has proposed an ODE model for predicting the growth profile of microalgae in response to medium concentration and light flux into culture to help the operators with cultivation planning. The proposed model has been built by extending the logistic equation in two steps based on the experimentally obtained growth profile of ACCB1808 (*Monoraphidium* sp.) under 16 conditions consisting of the combinations of incident light fluxes into culture and initial medium concentrations. Specifically, we have firstly constructed the MR model (Eq. (5)) that considers only the effect of light flux into culture, assuming that the medium concentration is high. In other words, the MR model captures the growth profile before reaching stationary phase under sufficient-high initial medium concentration conditions. Next, we have extended the MR model to incorporate the effect of the medium concentration in (Eq. (6)), where the Monod-type model was introduced based on the experimentally measured growth profile. The predictive ability of the proposed model has then been evaluated by a cross-validation method, and it has been shown that the predicted growth kinetics agrees with experimental data as shown in Fig. 2 (B). Finally, model-guided cultivation planning has been shown as a demonstration example, where the initial medium concentration and the incident light flux into culture have been planned for harvesting predefined target biomass at target time. The proposed model streamlines the planning process of cultivation cycles that satisfy various practical demands such as cost effectiveness and the schedule of operators.

## Data, codes and materials

The program codes used in this study are available at GitHub (https://github.com/hori-group/logistic_eq_for_cultivation_planning).

## Authors’ contributions

Y. Hirokawa and Y. Hori conceived the study. K.K, A.K, and Y. Hori developed the mathematical model, computational methods, and program codes. Y. Hirokawa performed cultivation experiments and data analysis. K.K., Y. Hirokawa, and Y. Hori drafted the manuscript. All authors gave feedback to the manuscript and gave final approval for publication.

## A Experimental conditions for cultivation

All ACCB1808 cultures were incubated in a space of which room temperature was controlled at 25°C with continuous aeration including 1.5 % carbon dioxide. LED light was used for growth, and light intensities were altered for incubation conditions. Photosynthetic photon flux density (PPFD) in each condition was measured by using Spectromaster C-7000 (SEKONIC; Tokyo, Japan). Optical density at 730 nm (OD), which is proportional to biomass concentration *N/V* where *N* is the biomass and *V* is the culture volume, was measured by using Spectrophotometer V-700 (JASCO; Tokyo, Japan). Biomass concentration *N/V* was counted by using microscope (Olympus; Tokyo, Japan) and a Thoma chamber.

For isolation and preculture, liquid and solid BG11 medium containing following components (per liter); 1.5 g of NaNO_3_, 31.4 mg of K_2_HPO_4_, 73.9 mg of MgSO_4_ · 7H_2_O, 36.8 mg of CaCl_2_ · 2H_2_O, 20.1 mg of Na_2_CO_3_, 1.12 mg of EDTA 2Na, 6.09 mg of citrate, 10.15 mg of ferric ammonium citrate, 1 mL of A6 solution, was used. A6 solution included following components (per liter); 2.86 g of H_3_BO_4_, 1.81 g of MnCl_2_ · 4H_2_O, 0.22 g of ZnSO_4_ · 7H_2_O, 0.39 g of Na_2_MoO_4_ · 2H_2_O, 0.079 g of CuSO_4_ · 5H_2_O, 0.049 g of Co(NO_3_)_2_ · 6H_2_O. To prepare solid medium, 15 g · L^−1^ agar was supplemented.

The growth profiles of ACCB1808 in different concentrations (from 73.9 to 739 mg · L^−1^) of MgSO_4_ · 7H_2_O in BG11 medium indicated that 6-fold concentration (443 mg · L^−1^) was appropriate (data not shown). Therefore, MgSO_4_ fortified medium was called modified BG11 medium and used for main culture cultivation.

In preculture cultivation, cells were inoculated into 60 mL of BG11 medium in a 100-mL test tube and incubated under continuous illumination (40 µE · m^−2^ · s^−1^) and aeration including 1.5 % carbon dioxide. Cells of preculture in late logarithmic growth phase were inoculated into 200 mL modified BG11 medium in 300-mL flask to an initial optical density at 730 nm (OD_0_) of 0.025.

## B Relation between incident light flux and its density

The incident light flux into culture *E*_in_ can be calculated by multiplication of incident light flux density *e*_in_ and illuminated area (2.826 × 10^−3^ m^2^):

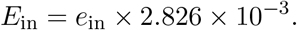

Assuming that the incident light linearly passes culture, the transmitted light flux through culture *E*_out_ is also calculated by

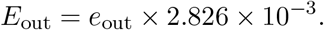

Based on Lambert-Beer’s law as mentioned in Appendix C, the light flux per cell *L*(*N* (*t*), *E*_in_) is obtained as Eq. (4) in Section 3.1.

## C Evaluation of extinction coefficient *K* based on LambertBeer’s law

The measured photosynthetic photon flux density (PPFD) under conditions with different culture depths *D* and optical densities at 730 nm (ODs) are summarized in Table 3. It should be noted that the unit of culture depth *D* is centimeter (cm) and that OD is proportional to the biomass concentration *N/V*, where *N* is the biomass and *V* is the culture volume. In Table 3, *e*_in_ and *e*_out_ correspond to PPFD in culture solution without cells and in culture with various ODs (OD=0.149–4.78), respectively.

**Table 3:** PPFD in different depth *D* and OD.

| $D$ (cm) \ OD | PPFD ($\mu\text{E} \cdot \text{s}^{-1} \cdot \text{m}^{-2}$) | | | | | | |
| --- | --- | --- | --- | --- | --- | --- | --- |
| | $e_{\text{in}}$ | $e_{\text{out}}$ | | | | | |
|  | 0 | 0.149 | 0.310 | 0.600 | 1.17 | 2.47 | 4.78 |
| 0 | 834.3 | 829.0 | 834.0 | 824.5 | 818.0 | 842.0 | 839.5 |
| 1 | 419.3 | 386.5 | 365.0 | 300.5 | 226.5 | 155.5 | 69.0 |
| 2 | 203.3 | 175.0 | 157.0 | 120.0 | 77.0 | 40.0 | 9.5 |
| 3 | 114.3 | 95.0 | 82.0 | 58.0 | 33.5 | 11.5 | 1.0 |
| 4 | 75.0 | 58.5 | 48.0 | 31.0 | 15.0 | 3.0 | N/A |
| 5 | 52.0 | 38.0 | 29.5 | 17.5 | 6.5 | 0.5 | N/A |
| 6 | 39.0 | 26.0 | 20.0 | 10.0 | 3.0 | N/A | N/A |

Optical density (OD) can be converted to biomass concentration *N/V* by

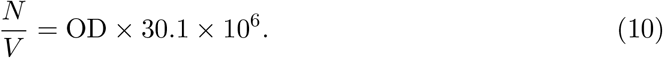

Thus, OD of 0.149, 0.310, 0.600, 1.17, 2.47 and 4.78 in Table 3 correspond to 4.55 × 10^6^, 10.1 × 10^6^, 17.6 × 10^6^, 34.9 × 10^6^, 69.8 × 10^6^ and 144 × 10^6^ cell · mL^−1^, respectively.

Figures 5 (A) and 5 (B) show relative logarithmic PPFD (= log(*e*_out_*/e*_in_)) for various culture depths *D* at particular OD (OD=0.149–4.78) and for various ODs at particular culture depth (1–6 cm), respectively. The relative logarithmic PPFD was negatively proportional to culture depth *D* and OD. This suggested that Lambert-Beer’s law was applicable to the culture solution. In other words,

**Figure 5:**
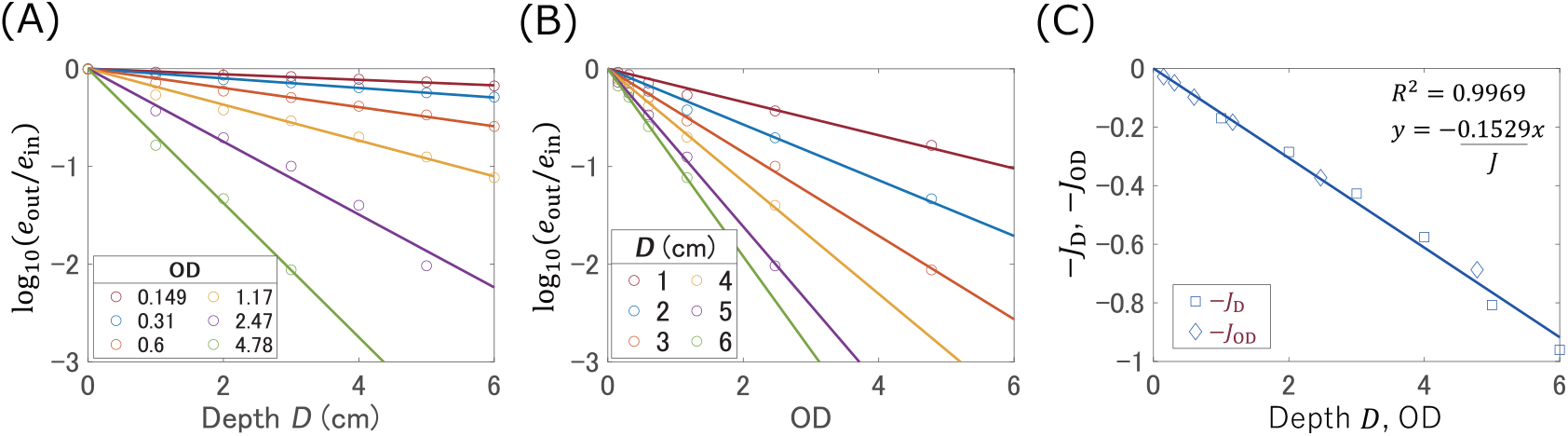
(A) Relationships between relative logarithmic PPFD (= log(*e*_out_*/e*_in_)) and culture depths *D* at particular OD (OD=0.149–4.78). Circles and solid lines show experimental data and the fitted line with slope (−*J*_OD_), respectively. (B) Relationships between relative logarithmic PPFD and ODs at particular culture depth (1–6 cm). Circles and solid lines show experimental data and the fitted line with slope (−*J*_D_), respectively. (C) Relation of −*J*_OD_ and −*J*_D_ with depth *D* and OD. Diamonds and squares show slopes *J*_OD_ and *J*_D_ obtained in panels (A) and (B), respectively. Solid line show slope of the fitted line for *J*.

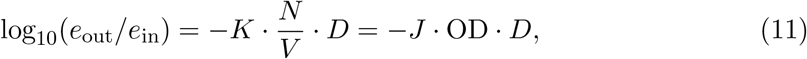

where *K* and *J* are the extinction coefficients defined for biomass concentration *N/V* and OD, respectively. The relationship between *K* and *J* can be written by *J* = *K* ×30.1×10^6^, according to Eq. (10). It should be noted that, by convention, the natural number *e* is also used as the base of logarithm in Eq. (11). In that case, the constants *K* and *J* should be redefined by the change of base formula.

To obtain the extinction coefficients, least square fittings were performed. Specifically, Eq. (11) is rewritten as

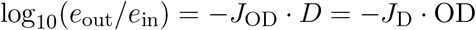

where *J*_OD_ is defined by *J*_OD_ := *J* ×OD and *J*_D_ is defined by *J*_D_ := *J* × *D*. Then, *J*_OD_ and *J*_D_ were calculated from the slopes of the fitted lines in Figs. 5 (A) and 5 (B), respectively. The extinction coefficient *J* was then obtained as *J* = 0.153 from the slope in Fig. 5 (C), which plotted *J*_OD_ and *J*_D_. The value of *J* = 0.153 corresponds to

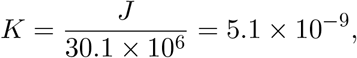

which is the extinction coefficient of ACCB1808 in the unit of mL · cm^−1^ · cell^−1^. The assessed extinction coefficient *K* = 5.1 × 10^−9^ mL · cm^−1^ · cell^−1^ was used for the prediction of the MR model in Section 3.2 and the full model in Section 4.2, and for the demonstration of cultivation planning in Section 5.

## References

[1] Saadaoui I, Rasheed R, Aguilar A, Cherif M, Al Jabri H, Sayadi S, et al. Microalgal-based feed: promising alternative feedstocks for livestock and poultry production. Journal of Animal Science and Biotechnology. 2021;12(1):1–15. doi: 10.1186/s40104-021-00593-z.

[2] Nagappan S, Das P, AbdulQuadir M, Thaher M, Khan S, Mahata C, et al. Potential of microalgae as a sustainable feed ingredient for aquaculture. Journal of Biotechnology. 2021;341:1–20. doi: 10.1016/j.jbiotec.2021.09.003.

[3] Ren Y, Sun H, Deng J, Huang J, Chen F. Carotenoid production from microalgae: biosynthesis, salinity responses and novel biotechnologies. Marine Drugs. 2021;19(12):713. doi: 10.3390/md19120713.

[4] Patel A, Karageorgou D, Rova E, Katapodis P, Rova U, Christakopoulos P, et al. An overview of potential oleaginous microorganisms and their role in biodiesel and omega-3 fatty acid-based industries. Microorganisms. 2020;8(3):434. doi: 10.3390/mi-croorganisms8030434.

[5] Yee W. Microalgae from the Selenastraceae as emerging candidates for biodiesel production: a mini review. World Journal of Microbiology and Biotechnology. 2016;32(4):1–11. doi: 10.1007/s11274-016-2023-6.

[6] Li S, Li F, Zhu X, Liao Q, Chang JS, Ho SH. Biohydrogen production from microalgae for environmental sustainability. Chemosphere. 2022;291:132717. doi: 10.1016/j.chemosphere.2021.132717.

[7] Yu X, Zhao P, He C, Li J, Tang X, Zhou J, et al. Isolation of a novel strain of Monoraphidium sp. and characterization of its potential application as biodiesel feedstock. Bioresource Technology. 2012;121:256–62. doi: 10.1016/j.biortech.2012.07.002.

[8] Zhao Y, Li D, Xu JW, Zhao P, Li T, Ma H, et al. Melatonin enhances lipid production in Monoraphidium sp. QLY-1 under nitrogen deficiency conditions via a multi-level mechanism. Bioresource Technology. 2018;259:46–53. doi: 10.1016/j.biortech.2018.03.014.

[9] Song X, Zhao Y, Han B, Li T, Zhao P, Xu JW, et al. Strigolactone mediates jasmonic acid-induced lipid production in microalga Monoraphidium sp. QLY-1 under nitrogen deficiency conditions. Bioresource Technology. 2020;306:123107. doi: 10.1016/j.biortech.2020.123107.

[10] Li X, Zhang X, Zhao Y, Yu X. Cross-talk between gama-aminobutyric acid and calcium ion regulates lipid biosynthesis in Monoraphidium sp. QLY-1 in response to combined treatment of fulvic acid and salinity stress. Bioresource Technology. 2020;315:123833. doi: 10.1016/j.biortech.2020.123833.

[11] Tale M, Ghosh S, Kapadnis B, Kale S. Isolation and characterization of microalgae for biodiesel production from Nisargruna biogas plant effluent. Bioresource technology. 2014;169:328–35. doi: 10.1016/j.biortech.2014.06.017.

[12] Mishra S, Mohanty K. Comprehensive characterization of microalgal isolates and lipid-extracted biomass as zero-waste bioenergy feedstock: an integrated biore-mediation and biorefinery approach. Bioresource technology. 2019;273:177–84. doi: 10.1016/j.biortech.2018.11.012.

[13] San Juan J, Mayol A, Sybingco E, Ubando AT, Culaba AB, Chen W, et al. A scheduling and planning algorithm for microalgal cultivation and harvesting for biofuel production. In: IOP Conference Series: Earth and Environmental Science. vol. 463. IOP Publishing; 2020. p. 012010.

[14] Ratkowsky D, Lowry R, McMeekin T, Stokes A, Chandler R. Model for bacterial culture growth rate throughout the entire biokinetic temperature range. Journal of bacteriology. 1983;154(3):1222–6. doi: 10.1128/jb.154.3.1222-1226.1983.

[15] Bernard O, Rémond B. Validation of a simple model accounting for light and temperature effect on microalgal growth. Bioresource technology. 2012;123:520–7. doi: 10.1016/j.biortech.2012.07.022.

[16] Bekirogullari M, Figueroa-Torres GM, Pittman JK, Theodoropoulos C. Models of microalgal cultivation for added-value products – a review. Biotechnology Advances. 2020;44:107609. doi: 10.1016/j.biotechadv.2020.107609.

[17] Béchet Q, Shilton A, Guieysse B. Modeling the effects of light and temperature on algae growth: state of the art and critical assessment for productivity prediction during outdoor cultivation. Biotechnology Advances. 2013;31(8):1648–63. doi: 10.1016/j.biotechadv.2013.08.014.

[18] Monod J. The growth of bacterial cultures. Annual review of microbiology. 1949;3(1):371–94. doi: 10.1146/annurev.mi.03.100149.002103.

[19] Ratkowsky DA, Olley J, McMeekin T, Ball A. Relationship between temperature and growth rate of bacterial cultures. Journal of bacteriology. 1982;149(1):1–5. doi: 10.1128/jb.149.1.1-5.1982.

[20] Morales M, Sánchez L, Revah S. The impact of environmental factors on carbon dioxide fixation by microalgae. FEMS microbiology letters. 2018;365(3):fnx262. doi: 10.1093/femsle/fnx262.

[21] Krebs CJ. Ecology: The experimental analysis of distribution and abundance. Pearson Education UK; 2013.

[22] Fujikawa H, Kai A, Morozumi S. A new logistic model for bacterial growth. Food Hygiene and Safety Science. 2003;44(3):155–60. doi: 10.3358/shokueishi.44.155.

[23] Fujikawa H, Kai A, Morozumi S. Improvement of new logistic model for bacterial growth. Food Hygiene and Safety Science. 2004;45(5):250–4. doi: 10.3358/shokueishi.45.250.

[24] Yang Z, Zhao Y, Liu Z, Liu C, Hu Z, Hou Y. A mathematical model of neutral lipid content in terms of initial nitrogen concentration and validation in Coelastrum sp. ha-1 and application in Chlorella sorokiniana. BioMed Research International. 2017;2017. doi: 10.1155/2017/9253020.

[25] Ram Y, Dellus-Gur E, Bibi M, Karkare K, Obolski U, Feldman MW, et al. Predicting microbial growth in a mixed culture from growth curve data. Proceedings of the National Academy of Sciences of the United States of America. 2019;116(29):14698–707. doi: 10.1073/pnas.1902217116.

[26] Peleg M, Corradini MG, Normand MD. The logistic (Verhulst) model for sigmoid microbial growth curves revisited. Food research international. 2007;40(7):808–18. doi: 10.1016/j.foodres.2007.01.012.

[27] Xu P. Analytical solution for a hybrid Logistic-Monod cell growth model in batch and continuous stirred tank reactor culture. Biotechnology and bioengineering. 2020;117(3):873–8. doi: 10.1002/bit.27230.

[28] Zarmi Y, Bel G, Aflalo C. Theoretical analysis of culture growth in flat-plate bioreactors: the essential role of timescales. Handbook of Microalgal Culture/Eds A Richmond, Q Hu Wiley-Blackwell. 2013:205–24.

[29] Poltronieri P, D’Urso OF. Biotransformation of agricultural waste and by-products. Food, Feed, Fibre, Fuel (4F) Economy. 2016:1–357.

[30] Masuda A, Horaguchi K, Kosaka S, Ozawa T, Kato M, Murakami K. The reduction of luminous transmittance in a microalgal culture tank with the increase of microalgal density and culture liquid thickness. Eco-engineering. 2006;18(1):3–8. doi: 10.11450/seitaikogaku.18.3 (in Japanese).

[31] James G, Witten D, Hastie T, Tibshirani R. An introduction to statistical learning. vol. 112. Springer; 2013.

