## Supplementary material for "A parametric logistic equation with light flux and medium concentration for cultivation planning of microalgae": Fig. S.1

### S. 1 Sequence alignment of 18S ribosomal DNA of ACCB1808

In this study, we originally isolated microalgae named ACCB1808. For phylogenetic analysis, 18S ribosomal DNA of ACCB1808 was sequenced. The analysis using Basic Local Alignment Search Tool (BLAST) showed that the sequence was 99.3%, 99.1%, and 99.1 % identical to that of *Monoraphidium* sp. LB59, *M. subclavatum* FBCC-A409, and *Monoraphidium* sp. HDMA-11, respectively, indicating that ACCB1808 was belonged to the genus of *Monoraphidium*. The sequence alignment of 18S ribosomal DNA was shown in Figure S.1.

*† These authors contributed equally to this work. Correspondence should be addressed to Yutaka Hori. Department of Applied Physics and Physico-Informatics, Keio University, 3-14-1, Yokohama, Kanagawa, 223-8522 Japan.

```
ACCB1808      1   TAGTCATATGCTTGTCTCAAAGATTAAGCCATGCATGTCTAAGTATAAACTGCTTATACTGTGAAACTGCGAATGGCTCATTAAATCAGTTATAGTTTAT   100

ACCB1808    101   TTGATGGTACCTCTACACGGATAACCGTAGTAATTCTAGAGCTAATACGTGCGTAAATCCCGACTTCTGGAAGGGACGTATTTATTAGATAAAAGGCCGA   200
LB59        101   ..........T.T...................A..................................................................   200
FBCC-A409   101   ..........T..........................................C..............................................   200
HDMA-11     101   ..........T.T...................A..................................................................   200

ACCB1808    201   CCGGGCTTTGCCCGACCCGCGGTGAATCATGATAACTTCACGAATCGCATAGCCTCGTGCTGGCGATGTTTCATTCAAATTTCTGCCCTATCAACTTTCG   300
LB59        201   .......C............................................................................................   300
FBCC-A409   201   .......C...............................................T............................................   300
HDMA-11     201   ....A.....T.........................................................................................   300

ACCB1808    301   ATGGTAGGATAGAGGCCTACCATGGTGGTAACGGGTGACGGAGGATTAGGGTTCGATTCCGGAGAGGGAGCCTGAGAAACGGCTACCACATCCAAGGAAG   400
LB59        301   ....................................................................................................   400
FBCC-A409   301   ....................................................................................................   400
HDMA-11     301   ....................................................................................................   400

ACCB1808    401   GCAGCAGGCGCGCAAATTACCCAATCCTGATACGGGGAGGTAGTGACAATAAATAACAATACCGGGCATTCAATGTCTGGTAATTGGAATGAGTACAATC   500
LB59        401   ....................................................................................................   500
FBCC-A409   401   ....................................................................................................   500
HDMA-11     401   ....................................................................................................   500

ACCB1808    501   TAAATCCCTTAACGAGGATCCATTGGAGGGCAAGTCTGGTGCCAGCAGCCGCGGTAATTCCAGCTCCAATAGCGTATATTTAAGTTGTTGCAGTTAAAAA   600
LB59        501   ....................................................................................................   600
FBCC-A409   501   ....................................................................................................   600
HDMA-11     501   ....................................................................................................   600

ACCB1808    601   GCTCGTAGTTGGATTTCGGGTGGGTTCCAGCGGTCCGCCTATGGTGAGTACTGCTGTGGCCCTCCTTTCTGTCGGGGACGGGCTCCTGGGCTTCACTGTC   700
LB59        601   ....................................................................................................   700
FBCC-A409   601   ............................................................................A.......................   700
HDMA-11     601   ....................................................................................................   700

ACCB1808    701   CGGGACTCGGATTCGACGATGATACTTTGAGTAAATTAGAGTGTTCAAAGCAAGCCTACGCTCTGAATACTTTAGCATGGAATATCGCGATAGGACTCTG   800
LB59        701   ....................................................................................................   800
FBCC-A409   701   ......CT..G.........................................................................................   800
HDMA-11     701   ....................................................................................................   800

ACCB1808    801   GCCTATCTCGTTGGTCTGTAGGACCGGAGTAATGATTAAGAGGGACAGTCGGGGGCATTCGTATTTCATTGTCAGAGGTGAAATTCTTGGATTTATGAAA   900
LB59        801   ....................................................................................................   900
FBCC-A409   801   ....................................................................................................   900
HDMA-11     801   ....................................................................................................   900

ACCB1808    901   GACGAACTACTGCGAAAGCATTTGCCAAGGATGTTTTCATTAATCAAGAACGAAAGTTGGGGGCTCGAAGACGATTAGATACCGTCGTAGTCTCAACCAT   1000
LB59        901   ....................................................................................................   1000
FBCC-A409   901   ....................................................................................................   1000
HDMA-11     901   ....................................................................................................   1000

ACCB1808   1001   AAACGATGCCGACTAGGGATTGGAGGATGTTCTTTTGATGACTTCTCCAGCACCTTATGAGAAATCAAAGTTTTTGGGTTCCGGGGGGAGTATGGTCGCA   1100
LB59       1001   ....................................................................................................   1100
FBCC-A409  1001   ....................................................................................................   1100
HDMA-11    1001   ....................................................................................................   1100

ACCB1808   1101   AGGCTGAAACTTAAAGGAATTGACGGAAGGGCACCACCAGGCGTGGAGCCTGCGGCTTAATTTGACTCAACACGGGAAAACTTACCAGGTCCAGACATAG   1200
LB59       1101   ....................................................................................................   1200
FBCC-A409  1101   ....................................................................................................   1200
HDMA-11    1101   ....................................................................................................   1200

ACCB1808   1201   TGAGGATTGACAGATTGAGAGCTCTTTCTTGATTCTATGGGTGGTGGTGCATGGCCGTTCTTAGTTGGTGGGTTGCCTTGTCAGGTTGATTCCGGTAACG   1300
LB59       1201   ....................................................................................................   1300
FBCC-A409  1201   ....................................................................................................   1300
HDMA-11    1201   ....................................................................................................   1300

ACCB1808   1301   AACGAGACCTCAGCCTGCTAAATAGTCACGTTCGCTTTTTGCGGATGGCCGACTTCTTAGAGGGACAGATGCTACAAAAGCATCGGAAGTATGAGGCAAT   1400
LB59       1301   .............................TGC............T.......................................................   1400
FBCC-A409  1301   ....................................................................................................   1400
HDMA-11    1301   .............................TGC.....C......T.......................................................   1400

ACCB1808   1401   AACAGGTCTGTGATGCCCTTAGATGTTCTGGGCCGCACGCGCGCTACACTGACGCATTCAACAAGCCTATCCTTGACCGAGAGGTCTGGGTAATCTTTGA   1500
LB59       1401   .......................................................G............................................   1500
FBCC-A409  1401   ....................................................................................................   1500
HDMA-11    1401   .....................................................A.G............................................   1500

ACCB1808   1501   AACTGCGTCGTGATGGGGATAGATTATTGCAATTATTAGTCTTCAACGAGGAATGCCTAGTAGGCGCATGTCATCAGCATGCGCCGATTACGTCCCTGCC   1600
LB59       1501   ...C................................................................A.........T.....................   1600
FBCC-A409  1501   ...................................................................GAT.......ATC....................   1600
HDMA-11    1501   ...C.T..............................................................A.........T.....................   1600
```

```
ACCB1808    1601  CTTTGTACACACCGCCCGTCGCTCCTACCGATTGGGTGTGCTGGTGAAGTGTTCGGATTGGCGAAGCGGGTGGCAACACTTGCTTTTGCCGAGAAGTTCA  1700
LB59        1601  ....................................................................................................  1700
FBCC-A409   1601  ..............................................................AG....................................  1700
HDMA-11     1601  ....................................................................................................  1700

ACCB1808    1701  TTAAACCCTCCCACCTAGAGGAAGGAGAAGTCGTAACAAGGTTTCC                                                        1746
LB59        1701  ..............................................                                                        1746
FBCC-A409   1701  ..............................................                                                        1746
HDMA-11     1701  ..............................................                                                        1746
```

Figure S.1: Multiple sequence alignment of 18S ribosomal DNA gene derived from originally isolated ACCB1808 and other Monoraphidium species deposited in GenBank. Accession number of LB59, FBCC-A409, and HDMA-11 are KP017571.1, MT169963.1, and MH340049.1, respectively.
